# Effect of landscape heterogeneity on diurnal raptor community richness and diversity in Jammu *Shiwaliks*, Jammu and Kashmir

**DOI:** 10.1101/2021.01.22.427758

**Authors:** Sudesh Kumar, Asha Sohil, Muzaffar Ahmed, Neeraj Sharma

## Abstract

In this study, we examined the richness and diversity of diurnal raptors along with their foraging guilds across different land uses in a sub-tropical landscape during December 2016 to November 2018. A total of 80 vantage points, 19 line transects and 36 road transects were sampled in 33 sites in six different habitat types in the study area where we recorded 3409 individuals of 29 diurnal raptors in 2 orders and 3 families. Significant variation in bird abundance was observed among different habitat types, farmlands being more specious followed by pure forests, water bodies and forest-farmland interfaces. Among the seasons, summers recorded higher abundance followed by winter, monsoon and post-monsoon. A low diversity value (H′=2.22) however was observed for the whole study area with mean monthly highest recorded during February (H′=2.44) and least during June (H′=1.85). Most of the raptors observed for their food types and foraging were predators (n=22) and rest were carrion feeders (n=22). Fourteen among all observed diurnal raptors were winter visitors and 13 residents with 9 reported globally threatened. A moderately high richness of diurnal raptors substantiate high conservation value of these habitats especially the forest patches and farmlands and thus calls for effective management strategies for the conservation and proliferation of raptors in sub-tropical areas of Jammu region.

## Introduction

Raptors are among the most dynamic avian species [1] characterized by unique morphological characteristics including strong talons, hooked beaks adaptation to tearing and/or piercing flesh etc. [2]. Because of their life history traits, relatively low population densities; large home ranges [3–5] and high trophic level, raptors are more sensitive to human disturbances [6–7], environmental contamination [3] and extinction [8] than other bird species. Raptors being the top-order predators, are considered excellent indicators of habitat quality [9] and play a potent role in structuring biological communities [10–11], besides, their population dynamics provide useful information of ecosystem status they inhabit [12–15].

High habitat heterogeneity and prey diversity promotes species richness and abundance among raptor communities [16]. Low population densities, slow turnover rates [8,17–18] and their more susceptibility to anthropogenic threats [6–7] leads to huge population decline [19]. Threats to raptors include habitat alteration [20–25], killings [26–27], poisoning [28–34], electrocution [35–38,], collisions with human made structures and vehicles [17,39–40], road killing [41], human consumption [42–43], climate change [44–49] and many more. Most of the raptors are seriously threatened all over the globe like Europe [50–51], Asia, Middle East and Africa [1,52–53]. The vultures across the Indian sub-continent have witnessed a catastrophic decline in their population following the introduction of diclofenac as a veterinary drug in the 1990s [28]. Despite their charisma and immense ecological significance, there is currently no systematic global synthesis of status, threats, or conservation for all raptors [54]. Raptors being the top predators are key taxa in conservation planning [55], regular monitoring of their population status is essential for proper management of natural ecosystems [56]. It thus becomes extremely essential to understand the status and distribution of raptors in different countries and regions [57].

Diurnal birds of prey are the predominant avian predators for natural ecosystems and are amongst the most susceptible species to the impacts of habitat transformations and perturbations [17]. Asia is home to 127 diurnal raptors (accipitriformes and falconiformes) contributing 40% of the total 317 diurnal raptors found all around the globe [58]. Heterogeneous geography, varied topography and great climatic variability makes northwestern Himalayas biodiversity rich region [59]. At the micro scale, 15 species of raptors were recorded from Baltistan, Pakistan occupied Jammu and Kashmir [60], 25 from south-eastern part of Jammu and Kashmir [61] and 13 species from the plains of Jammu city [62]. At regional level, [63] reported 47 species of raptors in three families, Accipitridae (39 species), Falconidae (7) and Pandionidae (1) from the erstwhile state of Jammu and Kashmir (which includes Jammu, Kashmir and Ladakh division). Forty three among these have been reported from Jammu division only. Despite high raptor diversity and species of conservation interest, the information on distribution and diversity of diurnal raptor communities is scanty and equivocal for the sub-tropical forests in the region. Forming one of the largest groups among the birds in the region [63], the raptors have not attracted the attention of bird ecologists from this part of the region, except for a breeding record by [64]. The current work aims to document diversity, richness, abundance, habitat guild and threat status of raptor communities in the Shiwalik region of UT of Jammu and Kashmir.

## Material and Methodology

### Study area

Organized surveys were carried out in six different habitats in a subtropical region of Jammu Shiwaliks lying at 32^0^27′ and 33^0^50′ N and 74^0^19′ and 75^0^20′ E between 317m to 1010m asl elevation and an area coverage of *c*. 5000 Km^2^ (Fig 1) during December 2016 to November 2018. The study area comprises of heterogeneous landscapes including southern alluvial plains, fallow lands, agricultural areas, urban built-up areas and the pine clad slopes near the Nandani Wildlife Sanctuary extending eastwards to Kathua region of Jammu division (Table 1, Fig 1). The vegetation comprises of subtropical scrub, broad-leaved associates interspersed with patches of Chirpine at higher elevations. The climate is generally dry sub-humid type and the whole year is divisible into four distinct seasons *i.e*., spring (February, March - mid-April), summer (mid-April-June), rainy (July-mid-September), and winter (October-January). The maximum summer temperatures range between 36°C and 42°C while the area receives the average annual precipitation of ~ 1000 mm. The surveys were conducted in 33 sampling sites among six distinct ecosystems intended to cover a range of diverse habitats with varied degree of disturbances. The sampling sites have been categorized as pure forests, forest-farmland interfaces, farmlands, urban built-up, greenbelt parks and urban avenues and water bodies (Table 1).

**Fig 1:**
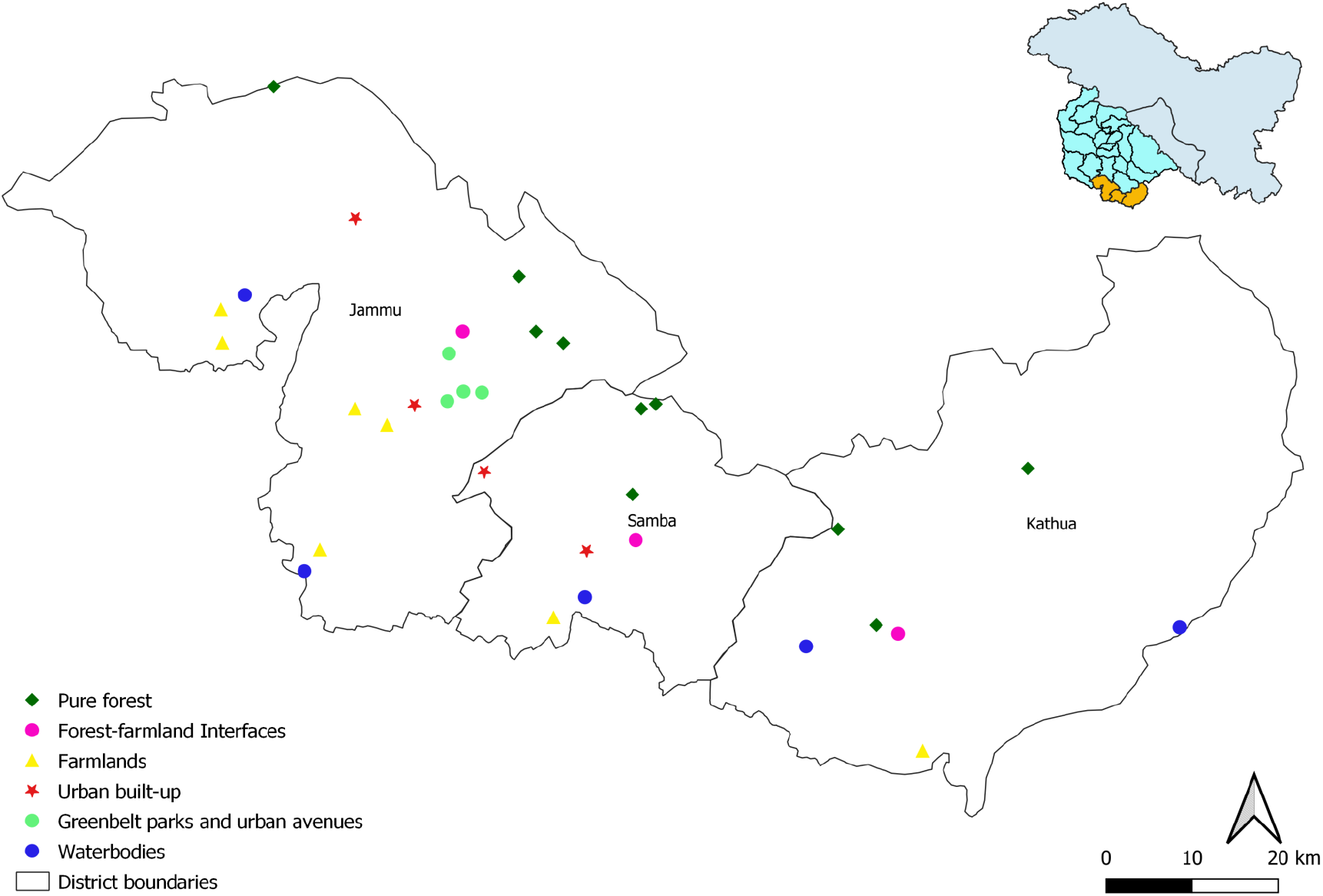
Map showing the location of sampling sites in the study area

**Table 1:**
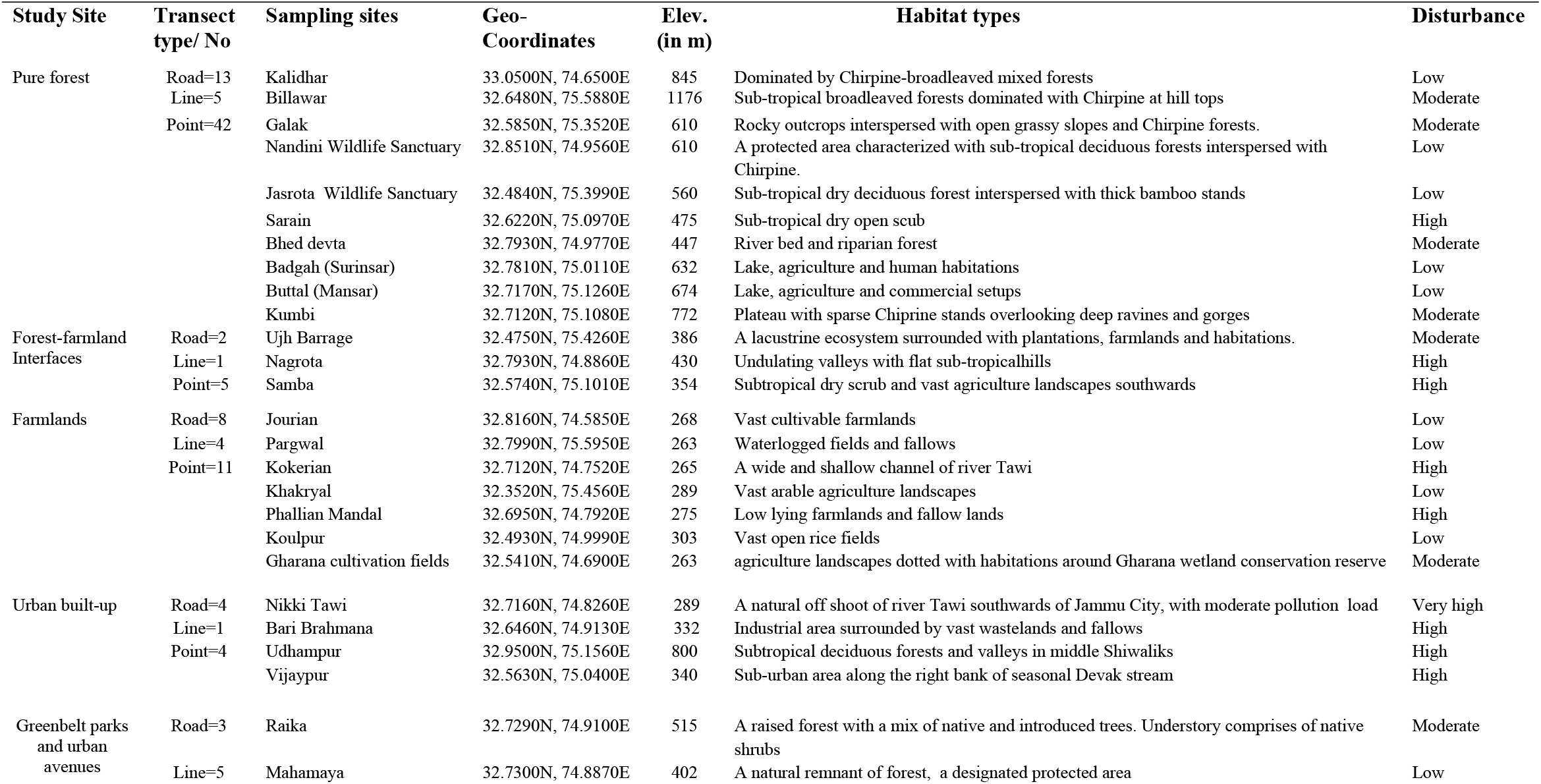
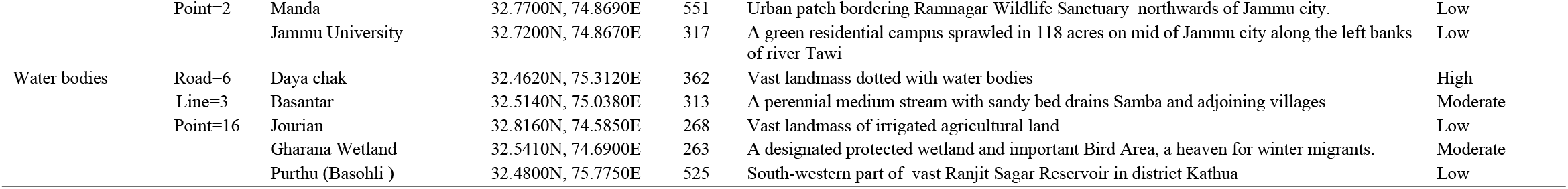
Characteristic features of the sampling sites with details on geo-features, transects and the level of disturbance

### Data Collection

The selection of sampling sites and sampling intensity was based on 15 days reconnaissance surveys conducted in the possible raptor occupancy sites in Jammu Shiwaliks. Transect method involving road transect of 25-60 km length [65], line transect (2-3 kms) and point count methods were used for counting the birds. For each transect, we recorded the number of species and individuals seen, activities observed and habitat occupied. To avoid the double count, transects were placed at 15 kms and 250 meters apart for road and line transects, respectively. The vantage points were fixed at the elevated areas for counting soaring raptors within circumference of 2-3 kms. A total of 80 vantage points, 19 line transects and 36 road transects were sampled in six different habitat types in the study area (Table 1).

Transects were walked / travelled during morning (2 hrs after sunrise) and evening hours (before dusk). Sampling was avoided during inclement weather conditions like storm, rain or fog. A few opportunistic sightings recorded close to the predefined sampling locations have also been included for the analysis. Field excursions were carried out with the aid of 10×50 binocular and Canon Eos 7D Mark II DSLR with 100*400 mm telephoto lens. Species identification was done by using standard field guide [66–68] and online bird identification platforms like J&K Birdlife, Indian Birds, Ask id’s of Indian Birds. For the systematic list of birds, the work of [69] was followed.

### Habitat Characterization and Foraging Behaviour

Based on their food composition and foraging behaviour, the raptors have been categorized as predators (hunt the prey, kill and consume) and carrion eaters (feed on dead animal matter) following [70]. Habitat affinities of raptors were also noted for six habitat types during the surveys. The species were classified in different threat categories following IUCN Red List of threatened species [71].

### Data Analysis

#### Richness and diversity attributes

Species diversity was considered the pooled number and summed abundance of each species across all months, seasons, study sites and entire study area. It was calculated using Shannon-Weiner [72] and [73] Simpson’s indices while species richness, taken as the number of species per unit area [74] was obtained by [75] and [76] index. Evenness was calculated following [77] and [78]. The statistical analysis was performed in PAST software package ver. 3.06 [79]. The Relative abundance (RA) of the birds was calculated by using the following formula;

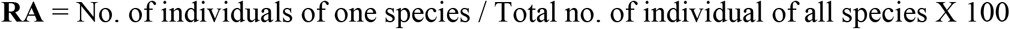

Significant differences in the species abundance among the four seasons *i.e*., Winter, Summer, Monsoon and Post-monsoon was compared using one-way ANOVA followed by Tukey’s multiple comparison tests [80]. Since the data was not normal (Shapiro Wilk W = 0.46, df = 116, p < 0.05), it was transformed to a logarithmic scale (log10) before the analysis. Similarly, Kruskal-Wallis test and Mann-Whitney U-test were applied to species abundance and richness values among the habitat types which appeared non-normal. The statistical tests were performed using SPSS (version 25) software packages with significance tested at p = 0.05.

## Results

### Species abundance

The study reported a total of 3409 individuals of 29 diurnal raptors in 2 orders and 3 families with 27 species belonging to family Accipitridae and one each to Pandionidae and Falconidae, respectively (Table 3). Most of the species have been observed in the farmlands (26 species) followed by pure forests (25 species), water bodies (22 species) and forest-farmland interfaces (16 species). The less specious habitats included urban built-up (11 species), green belt parks and urban avenues (9 species), each. Urban built-up recorded the highest mean population of 34.68 ± 21.44 raptors followed by pure forests (27.68 ± 9.71), farmlands (18 ± 4.81), forest-farmlands interfaces (13.58 ± 6.24), Green belt parks and urban avenues (11.48 ± 7.69, each) and water bodies (12 ± 3.44). Kruskal-Wallis test revealed a significant variation in bird abundance among the habitat types (H = 29.60; df = 5; p < 0.05) (Fig 2). The highest relative abundance was observed for *Milvus migrans lineatus* (RA=36.53) followed by *Neophron percnopterus* (RA=13.59) and *Aquila nipalensis* (RA=10.83). *Aquila fasciata* and *Aquila rapax* observed only twice during the entire survey period, showed the least relative abundance of 0.06, each (Table 3, Fig 3). When compared season wise, the highest species abundance was recorded during summer (40.93 ± 16.52) followed by winter (32.21 ± 11.19), monsoon (30.28 ± 13.58) and post-monsoon (14.13 ± 5.15). The seasons however did not play any significant role in governing the abundance of the species (ANOVA, F = 1.20, df = 3, p = 0.312) (Fig 4), the habitats exhibited, though. Pure forests showed a significant variation in raptor abundance when compared with forest-farmland interface (z = −2.10, n = 58, p = 0.035), urban built-up (z = −3.43, n = 58, p = 0.001) and green belt parks and urban avenues (z = −3.63, n = 58, p = 0.000); water bodies in relation to urban built-ups (z = −2.89, n = 57, p = 0.004); green belt parks and urban avenues (z = −3.16, n = 57, p = 0.002); farmlands with forest-farmland interfaces (z = −2.39, n = 58, p = 0.017), urban build-ups (z = −3.62, n = 58, p = 0.000) and green belt parks and urban avenues (z = −3.83, n = 58, p = 0.000). No-significant variation was however observed for the raptor abundance between water bodies and pure forests (z = −0.56, n = 57, p = 0.575), farmlands (z = −1.47, n = 57, p = 0.141) and forest-farmland interfaces (z = −0.74, n = 57, p = 0.456); urban built up with forest-farmland interfaces (z = −1.39, n = 58, p = 0.16) and green belt parks and urban avenues (z = −0.431, n = 58, p = 0.666).

**Fig 2.**
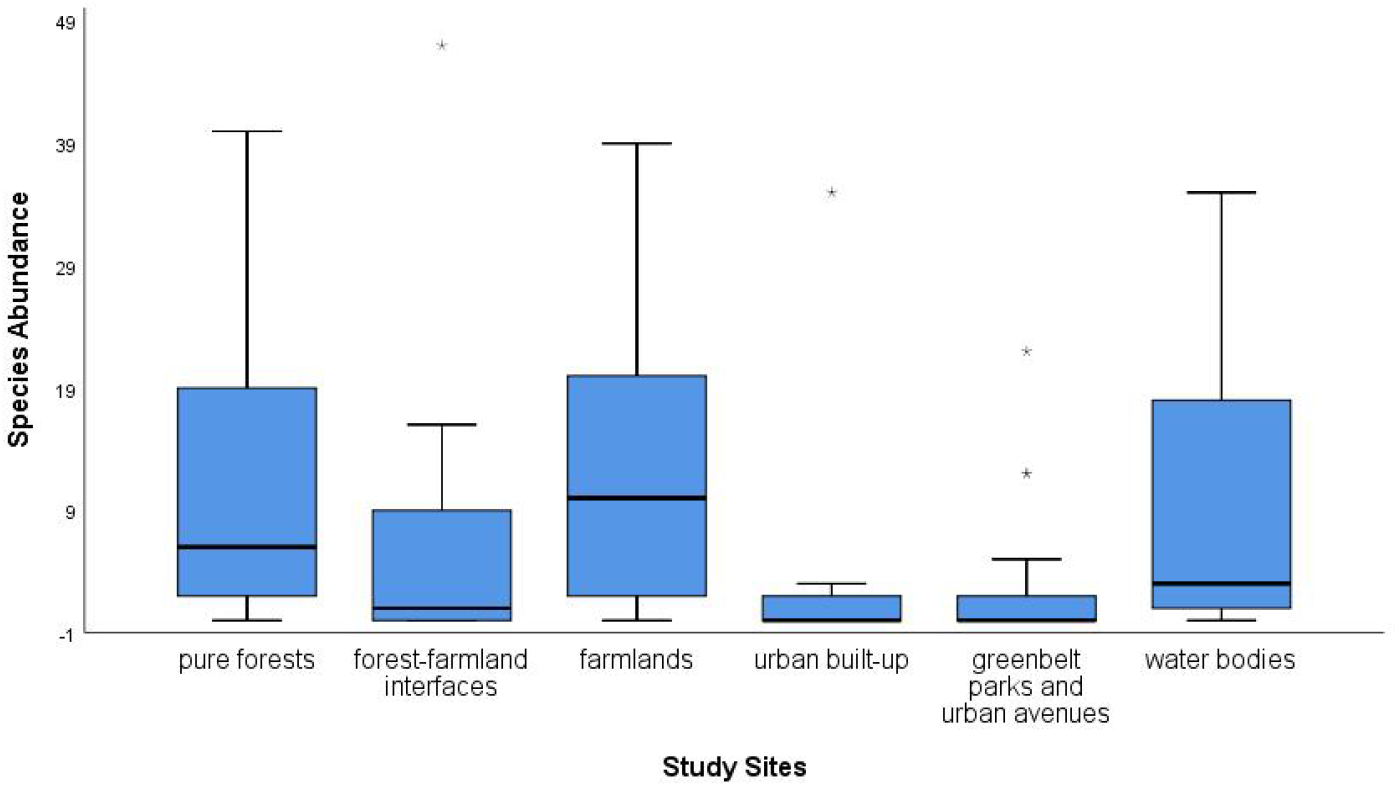
Bird species abundance among different habitats

**Fig 3:**
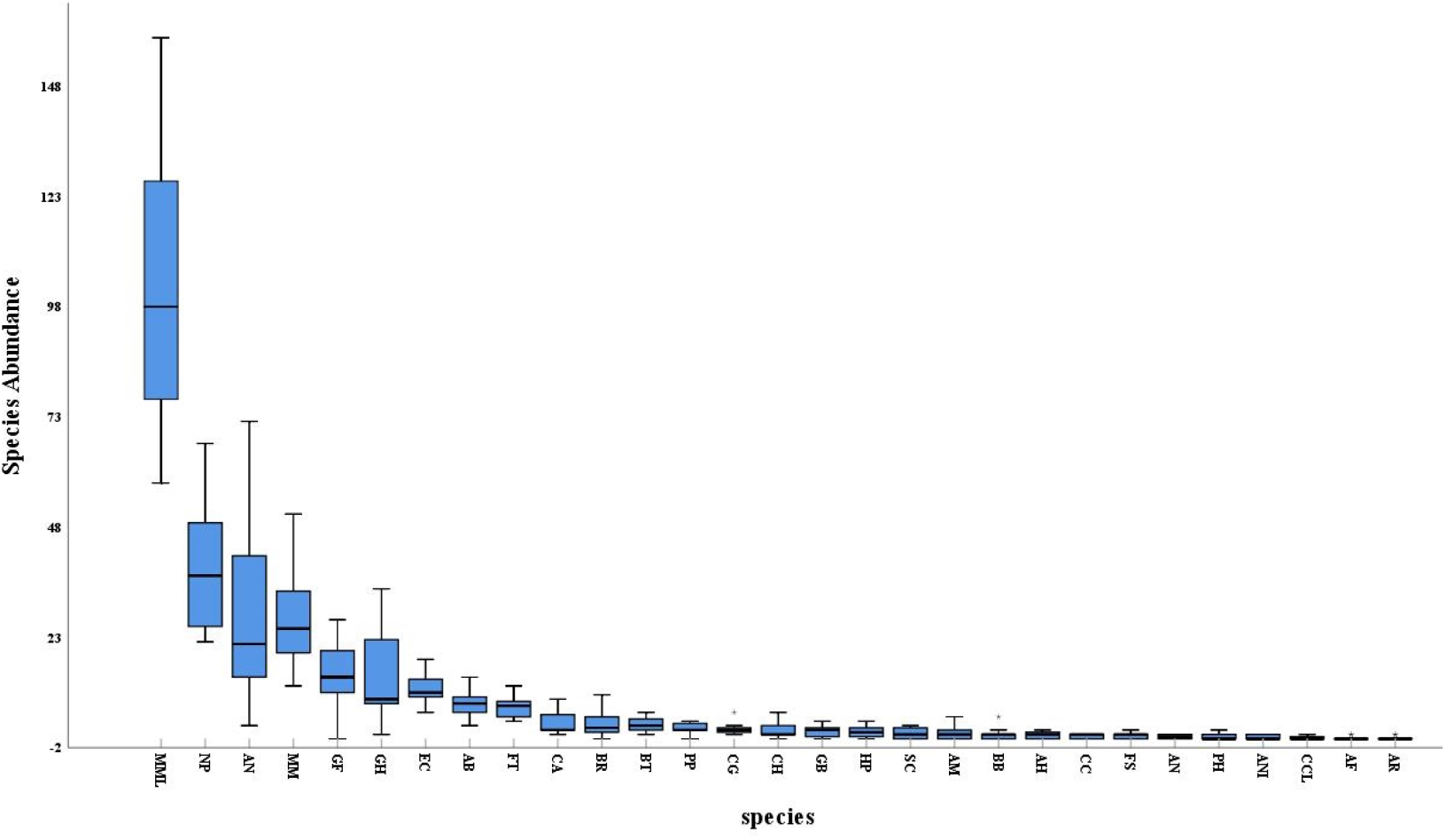
Mean abundance of each species MML: *Milvus migrans lineatus*; NP: *Neophron percnopterus*; AN: *Aquila nipalensis*, MM: *Milvus migrans*, GF: *Gyps fulvus*; GH: *Gyps himalayensis*; EC: *Elanus caeruleus*; AB: *Accipiter badius*; FT: *Falco tinnunculus*; CA: *Circus aeruginosus*; BR: *Buteo rufinus*; BT: *Butastur teesa*; PP: *Pernis ptilorhynchus*; CG: *Circaetus gallicus*; CH: *Clanga hastata*; GB: *Gyps bengalensis*; HP: *Hieraaetus pennatus*; SC: *Spilornis cheela*; AM: *Aegypius monachus*; BB: *Buteo buteo*; AH: *Aquila heliacal*; CC: *Circus cyaneus*; FS: *Falco subbuteo*; AN: *Accipiter nisus*; PH: *Pandion haliaetus*; ANI: *Accipiter nisus*; CCL: *Clanga clanga*; AF: *Aquila fasciata*; AR: *Aquila rapax*

**Fig 4:**
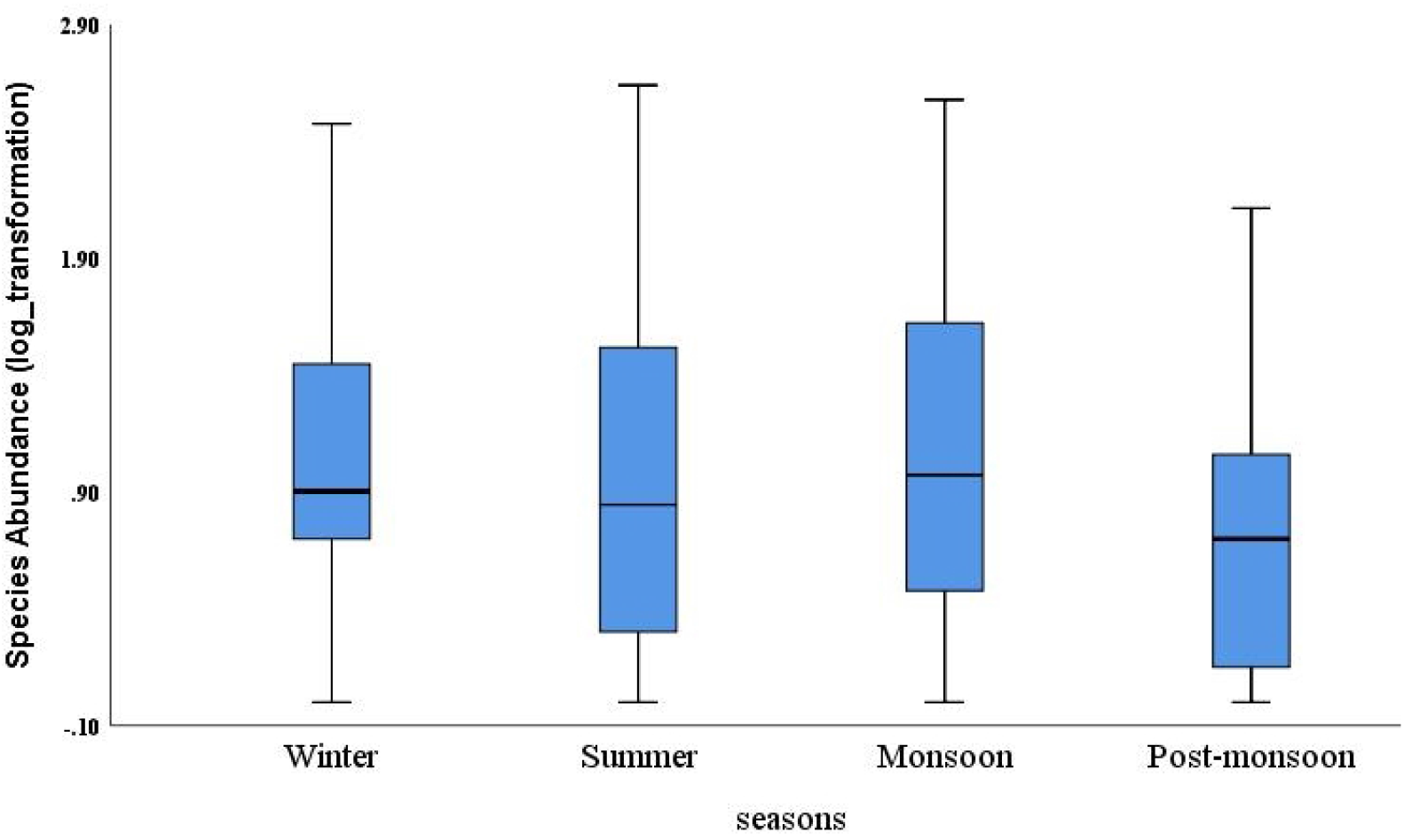
Bird species abundance among seasons

**Table 2:** Observed richness and diversity attributes of raptors across months and seasons

| **Month wise** | **Dec** | **Jan** | **Feb** | **W** | **Mar** | **Apr** | **May** | **S** | **Jun** | **Jul** | **Aug** | **Sep** | **M** | **Oct** | **Nov** | **PM** |
| --- | --- | --- | --- | --- | --- | --- | --- | --- | --- | --- | --- | --- | --- | --- | --- | --- |
| Taxa (S) | 27 | 25 | 24 | 29 | 25 | 22 | 19 | 27 | 16 | 18 | 16 | 21 | 22 | 21 | 25 | 27 |
| Individuals (N) | 282 | 323 | 329 | 934 | 474 | 392 | 321 | 1187 | 244 | 231 | 219 | 184 | 878 | 193 | 217 | 410 |
| **Diversity Index** |  |  |  |  |  |  |  |  |  |  |  |  |  |  |  |  |
| Simpson (1-D) | 0.80 | 0.82 | 0.87 | 0.85 | 0.85 | 0.78 | 0.74 | 0.81 | 0.71 | 0.80 | 0.78 | 0.75 | 0.77 | 0.83 | 0.82 | 0.83 |
| Shannon (H') | 2.24 | 2.28 | 2.44 | 2.41 | 2.25 | 2.01 | 1.87 | 2.11 | 1.85 | 2.09 | 2.01 | 1.94 | 2.02 | 2.23 | 2.29 | 2.31 |
| **Richness Index** |  |  |  |  |  |  |  |  |  |  |  |  |  |  |  |  |
| Menhinick (DMn) | 1.60 | 1.39 | 1.32 | 0.95 | 1.14 | 1.11 | 1.06 | 0.78 | 1.02 | 1.18 | 1.08 | 1.54 | 0.74 | 1.51 | 1.69 | 1.33 |
| Margalef (DMg) | 4.60 | 4.15 | 3.96 | 4.09 | 3.89 | 3.51 | 3.11 | 3.67 | 2.72 | 3.12 | 2.78 | 3.83 | 3.1 | 3.8 | 4.46 | 4.32 |
| **Species Estimate** |  |  |  |  |  |  |  |  |  |  |  |  |  |  |  |  |
| Equitability (J) | 0.68 | 0.70 | 0.77 | 0.71 | 0.70 | 0.65 | 0.63 | 0.64 | 0.67 | 0.72 | 0.72 | 0.63 | 0.65 | 0.73 | 0.71 | 0.70 |

Where, **W**–Winter; **S**–Summer; **M**–Monsoons; **PM**–Post-monsoon

**Table 3:**
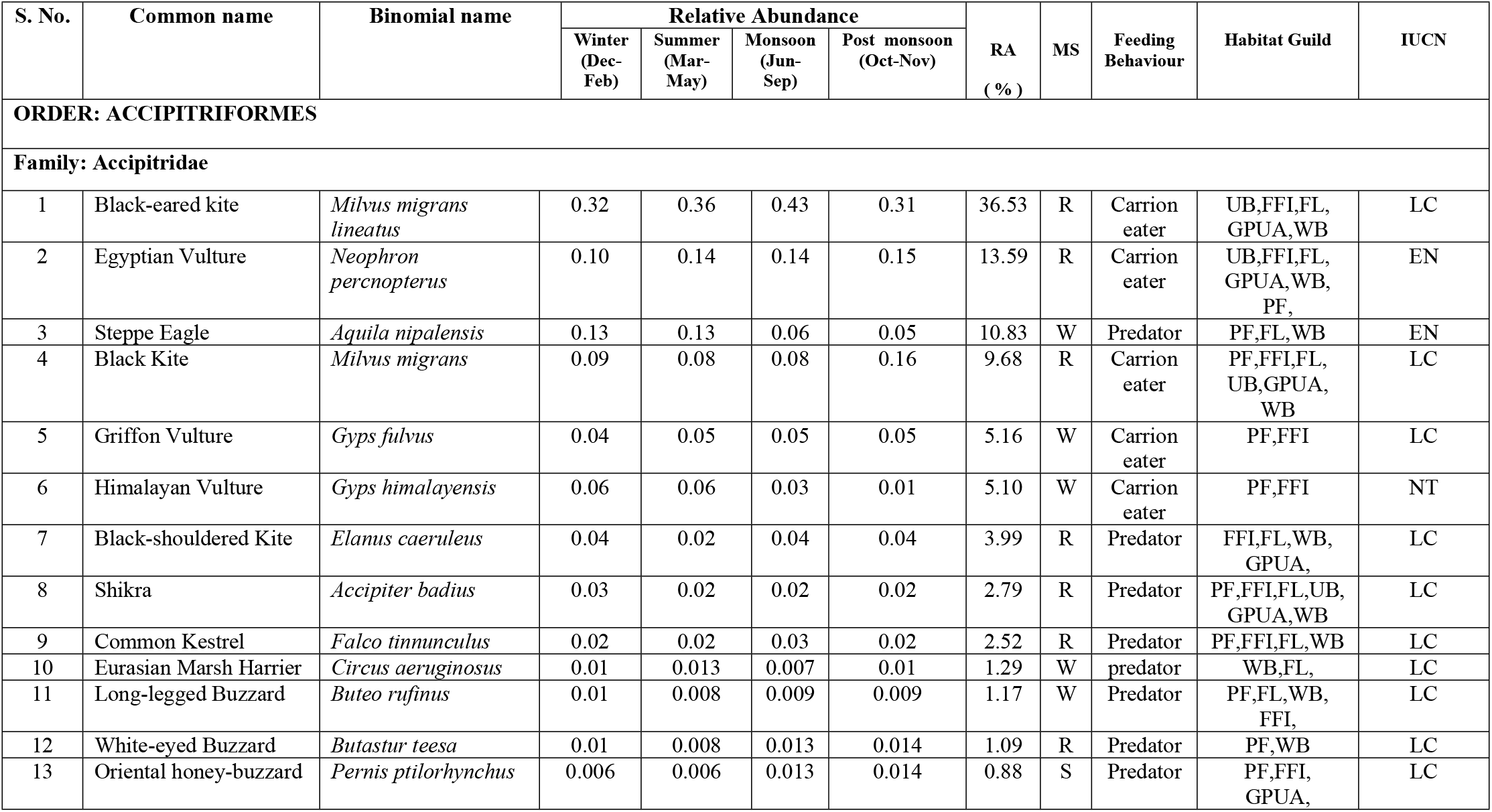
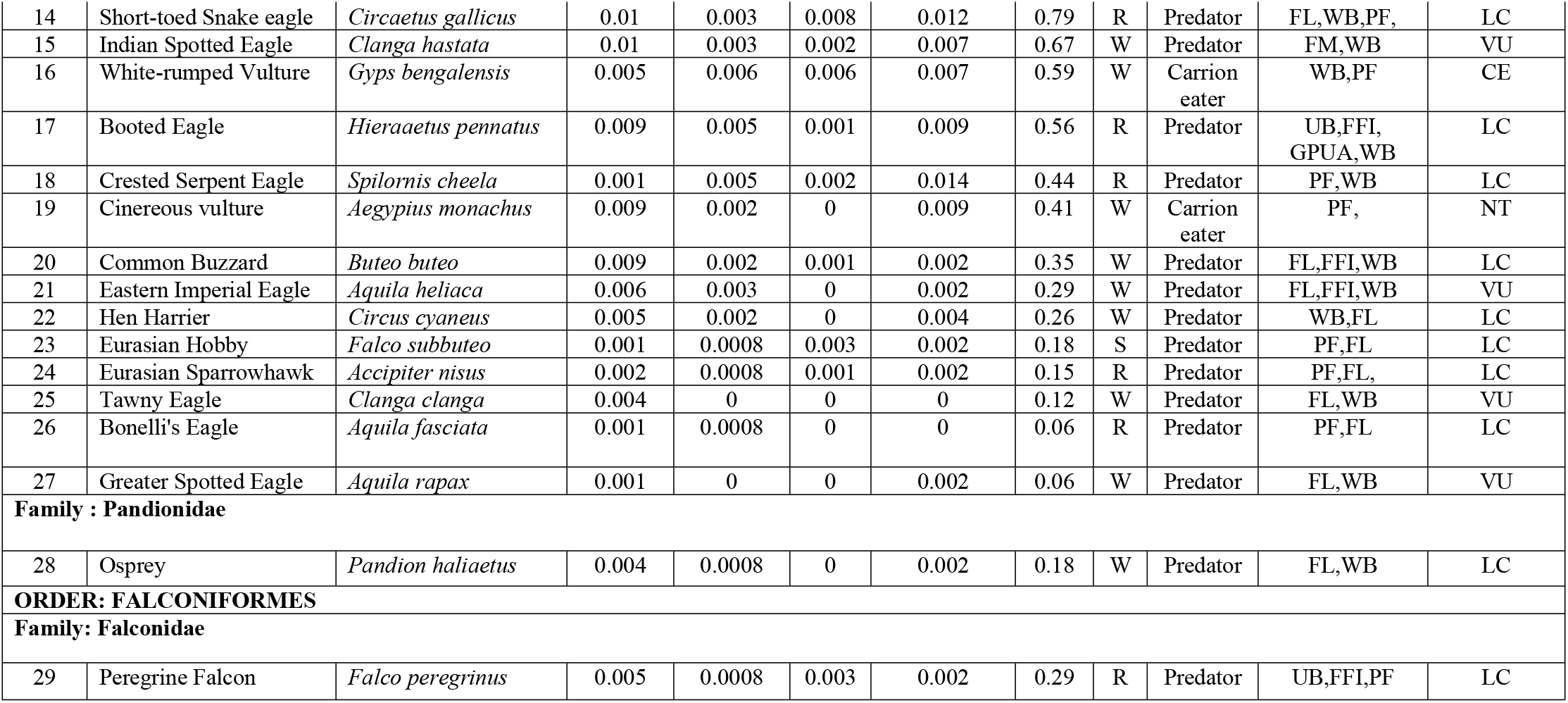
List of raptor species recorded in the study area along with their status, guilds and abundance

### Species richness and diversity

The richness and diversity attributes showed interesting results for different habitats. Maximum value for Menhinick’s and Margalef richness indices were recorded during November (D_Mn_= 1.69), and December (D_Mg_= 4.60), respectively (Table 2). When compared for the seasons, post-monsoon revealed maximum species richness (D_Mn_= 1.33; D_Mg_= 4.32) followed by winters (D_Mn_= 0.95; D_Mg_= 4.09), summers (D_Mn_= 0.78; D_Mg_= 3.67) while monsoon showed the least (D_Mn_= 0.74; D_Mg_= 3.1) (Table 2). Kruskal-Wallis test revealed a significant variation in bird species richness among the different habitats (H = 21.9; df = 5; p = 0.001). Post doc Mann-Whitney U-test showed variations in species richness for pure forests when compared with forest-farmland interfaces, urban build-ups, green belt parks and urban avenues (P < 0.05). *Neophron percnopterus, Milvus migrans* and *Accipiter badius* found in all the habitat types, were considered generalists. *Pandion haliaetus, Aquila rapax, Clanga clanga, Circus cyaneus, Aquila heliacal, Buteo buteo, Spilornis cheela, Circaetus gallicus, Butastur teesa, Circus aeruginosus, Neophron percnopterus* were mostly observed around the water bodies, dry riverbeds, sand beds and agricultural fields. The forest specialists included *Aquila nipalensis, Milvus migrans, Gyps fulvus, Gyps himalayensis, Falco tinnunculus, Buteo rufinus, Butastur teesa, Pernis ptilorhynchus, Gyps bengalensis, Falco subbuteo, Accipiter nisus* and *Aquila fasciata. Milvus migrans lineatus, Neophron percnopterus, Milvus migrans, Accipiter badius, Falco peregrines* dominated the urban landscapes. *Milvus migrans, Elanus caeruleus, Accipiter badius, Pernis ptilorhynchus, Hieraaetus pennatus* have been recorded around Green belt parks and urban avenues. Interestingly, the endangered *Neophron percnopteeus* and *Aquila nipalensis* revealed the high relative abundance in the study area.

The diurnal raptor communities showed a moderate diversity with the Shannon Weiner and Simpson’s diversity index values as 2.22 and 0.8, respectively. Mean monthly highest diversity (H′ = 2.44 and 1-D = 0.87), was observed during February (Table 2) and the least during June (H′= 1.85; 1-D = 0.71). Among the seasons, winter recorded the highest diversity (H′ = 2.41 and 1-D = 0.85) and monsoon the least (H′= 2.02; 1-D = 0.77) (Table 2). The species were observed evenly distributed during the winters with highest evenness index (J = 0.77) observed for February (Table 2). Among all, *Milvus migrans lineatus* and *Elanus caeruleus* showed the highest diversity (H′ = 2.44, 1-D = 0.90) whereas *Aquila fasciata* and *Aquila rapax* were least diverse (H′ = 0.69) each.

### Whittaker curves and Correspondence analysis ordination

The species diversity among different habitat types is well reflected by Whittaker curves (Fig 5) where urban built-ups and green-belt parks and urban avenues are ranked high with high raptor abundance followed by pure forests and forest-farmland interfaces whereas the farmlands and water bodies ranked low. Pure forests, water bodies, farmlands and forest-farmland interfaces with slanting curves reveal high species richness and evenness whereas the urban built-up and green belt parks and urban avenues behave vice-versa.

**Fig 5:**
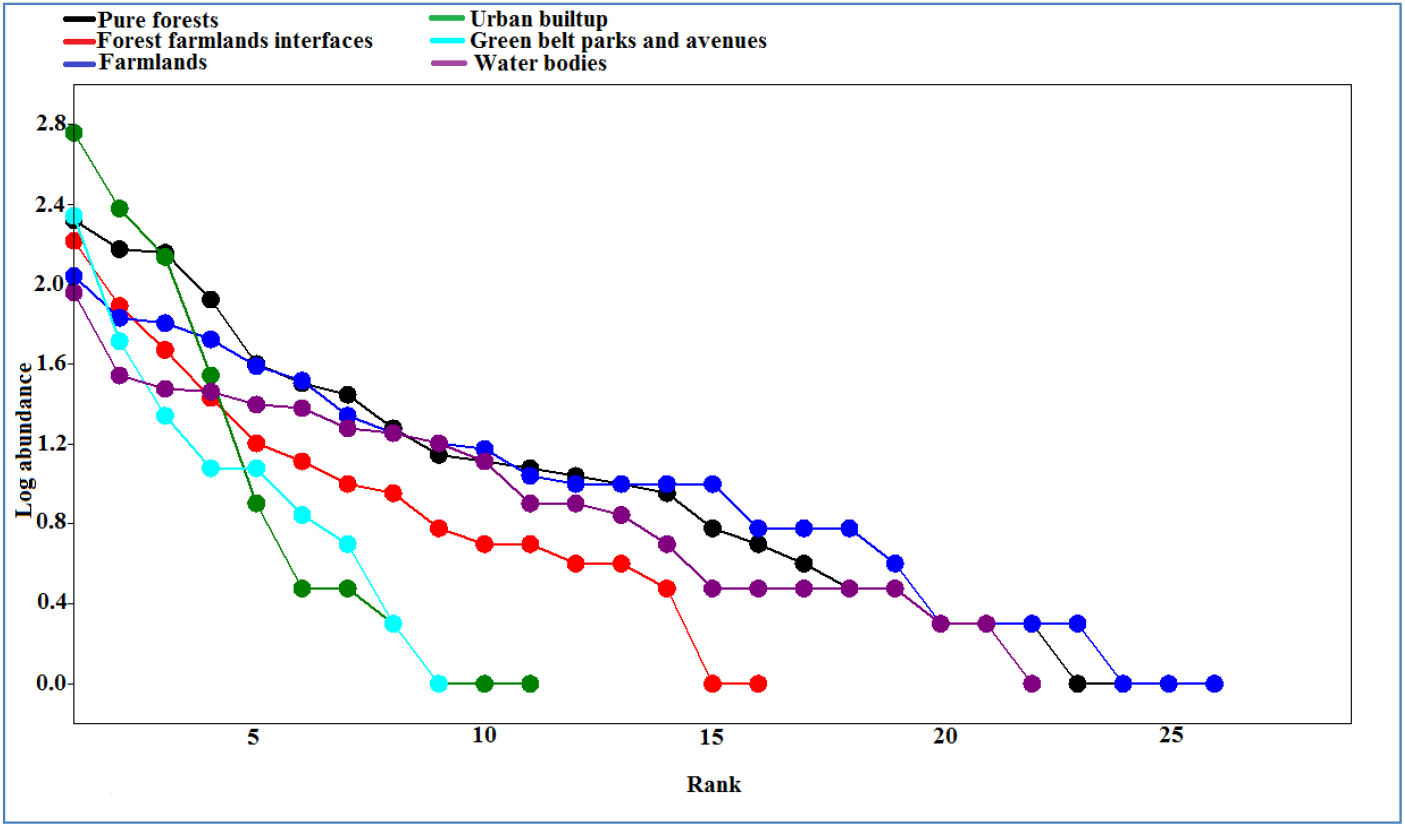
Diversity of species among the habitat types.

Correspondence analysis ordination plot reveals the bird community composition among the habitat types (Fig 6). Most of the species have their distribution restricted to pure forests, water-bodies and farmlands. The pure forests supported the habitats of Crested Serpent eagle (*Spilornis cheela*), Eurasian Hobby (*Falco subbuteo*), Himalayan Vulture (*Gyps himalayensis*), Griffon vulture (*Gyps fulvus*), Eurasian Sparrowhawk (*Accipiter nisus*) and Cinereous vulture (*Aegypius monachus*) whereas farmlands harbored Peregrine Falcon (*Falco peregrines*), Oriental honey Buzzard (*Pernis ptilorhynchus*), Booted Eagle (*Hieraaetus pennatus*), Bonelli’s Eagle (*Aquila fasciata*), Steppe eagle (*Aquila nipalensis*), Shikra (*Accipiter badius*), White-rumped vulture (*Gyps bengalensis*), Indian spotted Eagle (*Clanga hastata*), Greater Spotted Eagle (*Aquila rapax*) and Black shouldered Kite (*Elanus caeruleus*). Forest-farmlands and water bodies shared Kites, Eagles, Buzzards and Harriers whereas Kestrels, Osprey, Eastern Imperial Eagle (*Aquila heliacal*), Short-toed Snake Eagle (*Circaetus gallicus*) were found scattered around water bodies. The generalists like Egyptian vulture (*Neophron percnopterus*), Black kite (*Milvus migrans*) and Black-eared kite (*Milvus migrans lineatus*) occupied Urban built up and Green belt parks and avenues.

**Fig 6:**
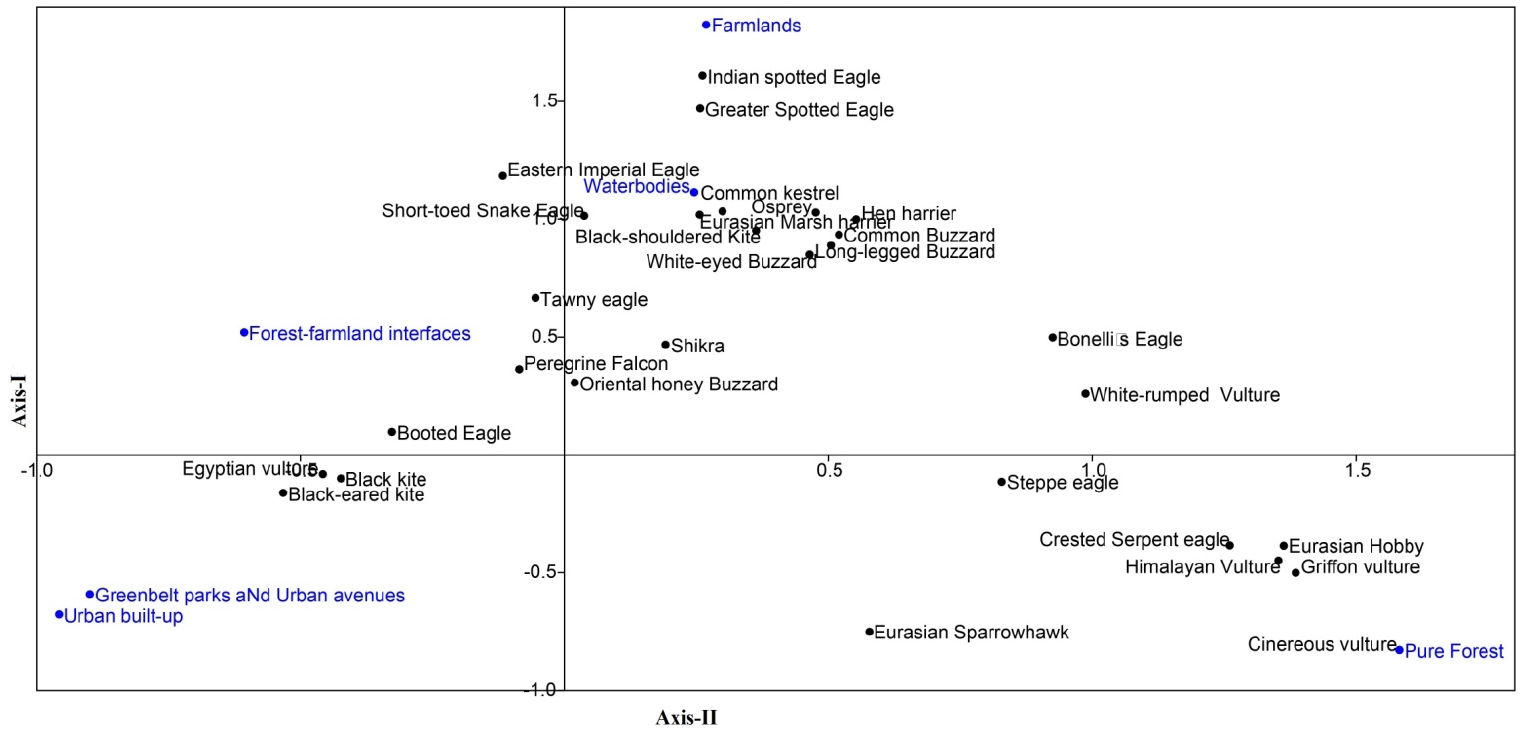
Correspondence analysis ordination between different habitats

### Feeding, migratory and conservation status

Our observation on food type and feeding behavior revealed that most of the raptors were predators (n= 22) and rest were carrion feeders (n= 7) (Table 3). Fourteen among the total species observed were winter visitors and these included *Aquila nipalensis, Gyps fulvus, Gyps himalayensis, Circus aeruginosus, Buteo rufinus, Clanga hastata, Gyps bengalensis, Aegypius monachus, Buteo buteo, Aquila heliacal, Circus cyaneus, Clanga clanga, Aquila rapax*and *Pandion haliaetus. Pernis ptilorhynchus* and *Falco subbuteo* were summer visitors whereas the remaining 13 were all residents (Table 3). Nine among the total species so observed were classified as globally threatened (IUCN, 2020). These included *Gyps bengalensis* (critically endangered); *Neophron percnopterus* and *Aquila nipalensis* (endangered), *Clanga hastata, Aquila heliacal, Clanga clanga* and *Aquila rapax* (vulnerable); *Gyps himalayensis* and *Aegypius monachus* (near threatened). Twenty species belonged to the least concern category (Table 3).

## Discussion

### Richness and Abundance

South and southeast Asia is home to high raptor diversity with 219 species recorded so far and of these 102 are found in India [58]. The present study reports 29 species accounting for 42% of the total diurnal raptors in India. This may be attributed to the heterogeneity of landscapes supporting an array of habitats comprising of intact forest patches, protected areas (Jasrota, Nandini, and Surinsar-Mansar Wildlife sanctuaries), rocky ridges, vast fallows and agricultural fields, water bodies (rivers, streams, and ponds), floodplains and urban habitats infused with green belt parks, urban forests and green corridors that provide favorable space for nesting, breeding, perching and roosting and thus high raptor richness and abundance [3,81–84]. Landscape attributes play an important role in determining avian richness and abundance [85–86] which is high in mosaic lands [87–89] limited by suitable breeding habitat and specific nest-site requirements [1,3]. Raptors which have large home ranges encompass a wide range of habitats [90]. Statistical analysis showed a contrasting deviation in species richness among different habitat types. The raptor abundance varied significantly for pure forest, farmlands and water bodies. Variation in bird species abundance could be due to their migratory behavior, food availability, habitat condition, and breeding season of the species [91–92]. Human disturbance is another important factor affecting the abundance and richness of birds [93]. The mean monthly highest values of Menhinick and Margalef richness indices observed for November and December may be attributed to the winter migration. This leads to the overall population inflation and thus the increased diversity as well during the winters.

Presence of raptors in urban areas can be related to stable or abundant prey bases, novel environments, reduced competition, and additional nesting structures [94]. Same holds good with two most abundant raptors observed in the urban environment which included *Milvus migrans lineatus* and *Neophron percnopterus*. These generalist species exploit buildings, other human structures and ornamental plantations for shelter, nesting, and food sources (including human-subsidized foods) reaching highest densities in urban areas [95]. Perching sites close to roads (power lines and telephone poles) and road kills increase attractiveness of raptors to urban areas [96]. Urbanization increases biological homogenization and consequently urban adaptable species become increasingly widespread and locally abundant [97]. The green belt parks, avenues, urban forests and green corridors in the present study area act as an ecotonal zone and have been quite successful in providing refuge to the raptor species like *Milvus migrans lineatus, Neophron percnopterus, Milvus migrans, Elanus caeruleus, Accipiter badius, Pernis ptilorhynchus* and *Hieraaetus pennatus*. Rich plant diversity, availability of perching sites (trees, electricity poles, and telephonic poles) and food resources could be the reason for good species number of raptor in green belt parks and urban avenues.

Urban avoiders like eagles, hawks, and falcons [98] with specific habitat requirements [95] are more abundant in less-disturbed and natural habitats [95,99–100] that provide safe refugee, hostile environment and the prey species [90]. It comprises the species adapted to live in the interior of forests, migrants, nesting birds sensitive to human presence. The forest dwellers observed during this study included *Accipiter badius, Falco subbuteo, Accipiter nisus* and *Falco peregrines, Gyps fulvus, Gyps himalayensis, Buteo rufinus, Butastur teesa, Pernis ptilorhynchus, Circaetus gallicus, Spilornis cheela* and *Aegypius monachus*. Agricultural fields and large open spaces provide breeding and foraging habitats for many open space foragers such as the Eurasian buzzards (*Buteo buteo*), harrier species (*Circus* ssp.), etc [101–104]. Besides providing new habitats, irrigated crops provide higher food availability to birds of prey in the form of small mammals, voles and rodents which are ideal foods for western marsh-harrier [105–106], black kites [107–108], black-winged kites and some migrant raptors like booted eagle and steppe eagle [13,109–110]. High raptor abundance in the farmlands and forest farmland interfaces in the study area is attributed to the availability of perching sites (more artificial in the form of poles and high tension / mobile towers) and diverse food options.

### Feeding, Migratory and Conservation status

Food supply is an important factor governing the raptor density [3] as birds and mammals form the main food for raptors [90] besides reptiles, amphibians, fish and arthropods [111]. Pertinently the number of the carrion feeders among the total appeared low but three among seven carrion feeders, *Milvus migrans lineatus, Neophron percnopterus* and *Milvus migrans* were among the top four highly abundant species recorded during the study. Most of them were reported mainly from urban habitats along the road side, near the water bodies and at perches either eating or roosting. Remaining four carrion feeders i.e., *Gyps fulvus, Gyps himalayensis, Gyps bengalensis* and *Aegypius monachus* were reported from forests and/or forest farmlands interfaces feeding on cattle that die naturally attracting the large numbers of vultures. Carcass of large animals; deer, hares in forest are the potential source of carrion forming a potent food base for raptors in forests [90,112]. Unlike scavenging raptors, predatory raptors search for their prey visually and hunt those [113]. The observed richness of predatory raptors was quite high than the scavenging ones. This may be attributed to the availability of diverse food availability in mosaic habitats in the study area. [114] reported that food availability is the most important criteria for selecting suitable stopover sites for migratory species and large concentrations of raptors are common in wintering grounds with abundant prey [115] that holds good with the current study. Predatory winter migrants and resident raptors, *Aquila nipalensis, Elanus caeruleus, Circus aeruginosus, Buteo rufinus, Butastur teesa, Circaetus gallicus, Clanga hastata, Hieraaetus pennatus, Spilornis cheela, Accipiter badius, Buteo buteo Aquila heliacal, Clanga clanga, Aquila rapax, Pandion haliaetus* observed in the mosaic habitats, were commonly observed feeding on lizards, rodents, insects, birds, etc.

The migratory behaviour of different raptor species in the study area may be due to seasonal movement patterns, local and regional habitat changes, large-scale population changes, and climatic conditions [116–118]. Most of the winter migrants during the study were reported from paddy fields and other farmlands situated close to wetlands or open areas near streams or floodplains and few of them were recorded from forest areas during winters. Migratory species often use paddy fields in winter as foraging sites [119–120]. The forest also serve as alternative habitat for the migratory species which could use the area for resting, foraging till the return of favorable condition [121–122]. The occurrence of 16 migratory species of raptors supports the fact that study area is an important conservation priority area especially for the winter migrants. Out of 26 globally threatened raptor species in India [58] (Bird life international, 2017) nine were observed during the current surveys. *Neophron percnopterus* is resident to the study area and is second most abundant species recorded from the study area whereas *Aquila nipalensis, Gyps himalayensis, Clanga hastata, Gyps bengalensis, Aegypius monachus, Aquila heliacal, Clanga clanga* and *Aquila rapax* are winter migrant to the region.

### Conclusions

Our study highlights the importance of urban and suburban areas, forests, streams, floodplains and farmlands for raptor conservation. Combination of natural, semi natural, and urban habitats, are becoming hot spots of landscape diversity that allow a high number of species with contrasting habitat needs. The farmlands and forests harbored the highest species of raptors. Water bodies like streams, rivers and adjoining floodplains too attracted unique raptor species. The conservation value of these mosaic habitat types is not only related to the number of raptor species but also due to inclusion of raptors that need special conservation measures. Indeed, several threatened birds of prey reported from these habitats are rare or even absent in the adjacent protected areas of Jammu Shiwaliks. Holistic actions are needed to implement focusing on ecosystems and habitats essential for maintaining biological diversity, preventing species extinction and increasing the ecological scope for recovery of endangered species. This research work has great implication for combined effort in conservation of raptors in future especially from Himalayan region.

## Acknowledgments

The authors gratefully acknowledge Department of Wildlife Protection, Govt. of Union Territory of Jammu and Kashmir for granting necessary permits and providing logistic support during the surveys Rector, Bhaderwah Campus, University of Jammu is also thanked for providing the administrative support during this study. The help rendered by Mr. Dinesh Singh, Anil Thakar and Ajaz Quereshi, in conducted field surveys is highly appreciated and duly acknowledged

